# Identifying cancer pathway dysregulations using differential causal effects

**DOI:** 10.1101/2021.05.20.444965

**Authors:** Kim Philipp Jablonski, Martin Pirkl, Domagoj Ćevid, Peter Bühlmann, Niko Beerenwinkel

**Author notes:** To whom correspondence should be addressed **Corresponding authors** Correspondence to Niko Beerenwinkel. Equal contribution (Kim Philipp Jablonski, Martin Pirkl).

## Abstract

Signaling pathways control cellular behavior. Dysregulated pathways, for example due to mutations that cause genes and proteins to be expressed abnormally, can lead to diseases, such as cancer. We introduce a novel computational approach, called Differential Causal Effects (*dce*), which compares normal to cancerous cells using the statistical framework of causality. The method allows to detect individual edges in a signaling pathway that are dysregulated in cancer cells, while accounting for confounding. Hence, artificial signals from, for example, batch effects have less influence on the result and *dce* has a higher chance to detect the biological signals. We show that *dce* outperforms competing methods on synthetic data sets and on CRISPR knockout screens. In an exploratory analysis on breast cancer data from TCGA, we recover known and discover new genes involved in breast cancer progression.

## 1 Introduction

The complexity of cancer makes finding reliable diagnosis and treatment options a difficult task. Decades of research made the intractable disease better understood. However, many challenges remain due to its high variability and context specificity, e.g., regarding tissue and cell type [1]. Patients with common cancer types in early stages show promising survival rates, even though rare subtypes still show low survival rates due to different traits like a more aggressive disease progression [2–4].

It has been hypothesized that cancer diversity can at least in part be explained by heterogeneous mutational patterns. These patterns influence the activity of biological pathways at the cellular level [5, 6]. For example, signaling pathways consist of several genes, which regulate certain cell programs, such as growth or apoptosis. The programs are driven by the causal interaction of the genes, e.g., the up-regulation of one causes the up-regulation of another gene. The causal effect (CE) determines the strength of this causal interaction, e.g., by increasing the expression of gene *X* two-fold, the expression of its child *Y* increases four-fold. Thus, *X* has a causal effect on *Y* of 2 [7]. Understanding how these causal networks are perturbed in tumors is necessary for prioritizing drug targets, understanding inter-patient heterogeneity, and detecting driver mutations [8].

Traditionally, perturbed pathways are detected by assessing whether differentially expressed genes are members of the respective pathway more often than expected by chance. More sophisticated methods measure whether genes belonging to a pathway are localized at certain positions of a rank-ordered set of differentially expressed genes [9]. In such cases, a pathway is interpreted as a simple set of genes and all topological information concerning the functional interconnectivity of genes is ignored. It has been recognized that interactions among genes can have a significant effect on the computation of pathway enrichments. Some tools consider, for example, gene expression correlations to account for confounding effects and control the type I error while retaining good statistical power [10]. The underlying structure of gene interactions can thus be either estimated from the data used for the enrichment analysis, or obtained from existing databases. Canonical pathway databases such as the Kyoto Encyclopedia of Genes and Genomes (KEGG) [11] can then be incorporated as prior knowledge to guide the enrichment analysis using topological information of gene connectivity [12–14].

While such enrichment methods go beyond treating pathways as plain gene sets and incorporate topological information of molecular interactions, they often only report a global pathway dysregulation score [14]. An exception is PARADIGM, which records an inferred activity for each entity in the pathway under consideration for a given patient sample [15]. It does, however, not model causal effects, but only quantifies whether there is some general association among the genes like correlation. Differential causal effects (DCEs) on biological pathways have already been investigated in a formal setting [16–18], where a DCE is modeled as the difference between CEs for the same edge under two conditions. These methods infer the gene network from observational data, which is a difficult task due to the combination of typically low sample size and noise of real data. An incorrect network can result in biased estimation of CEs and DCEs. Additionally, none of these methods make use of the DCEs to compute a pathway enrichment score.

Here, we separate the problem of estimating the causal network and the CEs by replacing the former with the addition of prior knowledge in the form of biological pathways readily available in public databases [11, 19–22]. We make use of the general concept of causal effects in order to define differential CEs. Specifically, we estimate the CE of gene *X* on gene *Y* in normal samples and cancer samples and define the DCE as their difference. In particular, we compare the causal effects between two conditions, such as a malignant tissue from a tumor and a healthy tissue, to detect differences in the gene interactions. We propose Differential Causal Effects (*dce*), a new method which computes the DCE for every edge (i.e., molecular interaction) of a pathway for two given conditions based on gene expression data (fig. 1).

**Figure 1:**
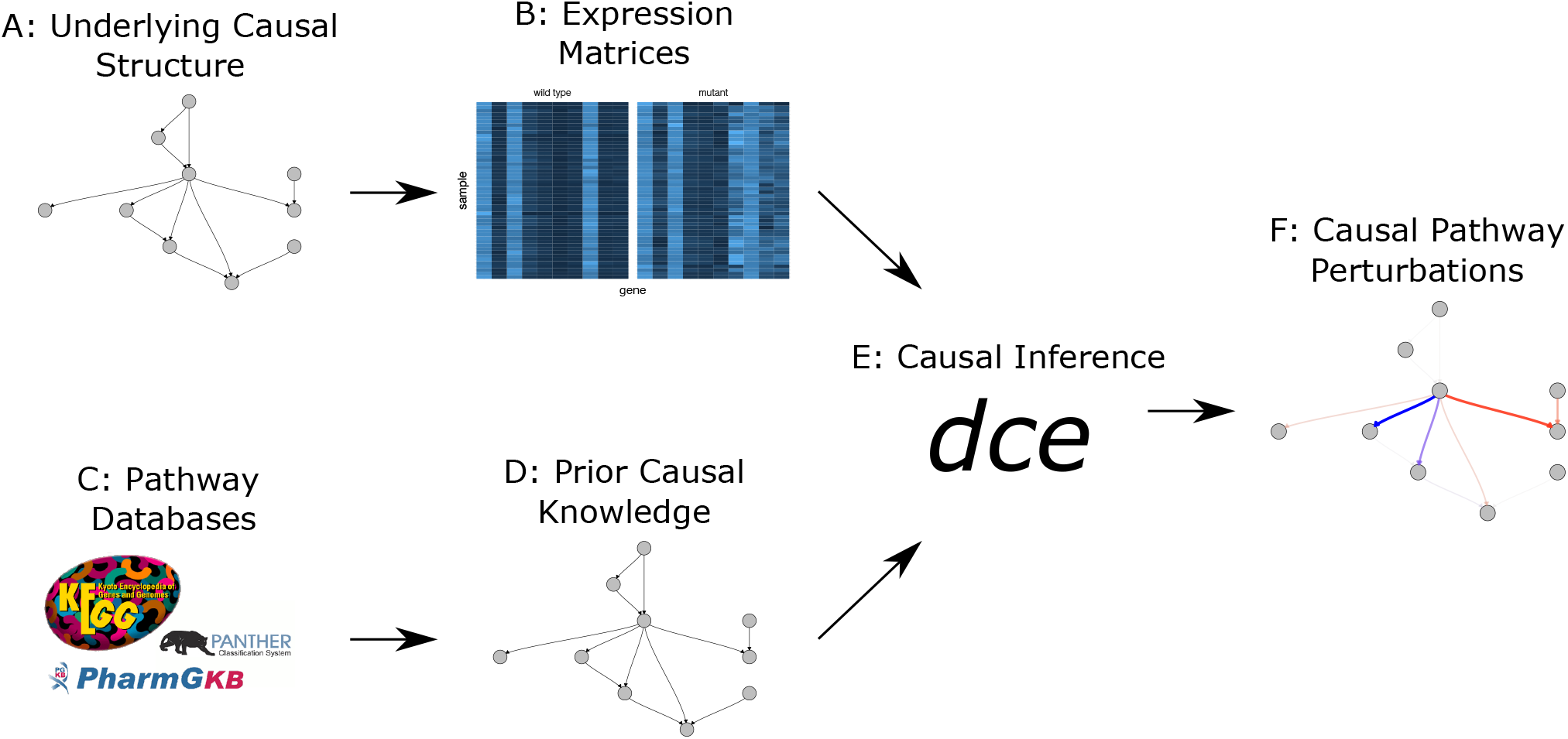
A causal network of genetic interactions in a biological pathway (A) is responsible for the observed wild type expression levels in a cell (B: wild type). A disease can lead to perturbations of these pathways and in turn generate altered expression levels (B: mutant). Pathway databases such as KEGG [11], PharmGKB [20] and Panther [21] curate genetic interaction data (C) and thus provide networks of putative causal interactions (D). Given the observed wild type and disease expression levels as well as the causal structure, *dce* fits a GLM for each edge to estimate causal effects (E). These correspond to causal perturbations (i.e., differential causal effects), e.g., an increase of causal effect strength from wild type to mutant is marked in blue. Negative differential causal effects are marked in red. The transparency of an edge corresponds to the magnitude of the associated effect) of the biological pathway caused by the disease and are important for diagnosis and treatment design (F).

This allows us to identify pathway perturbations at the individual edge level while controlling for confounding factors using the statistical framework of causality. By including the additional covariates constructed from the principal components of the design matrix, we also provide a methodological extension of our method to handle potential confounding that is unobserved in our data. For example, batch effects from different experimental laboratories are not necessarily known, but accounted for automatically. Our approach allows for computing pathway enrichments in order to rank all networks in large pathway databases to identify cancer specific dysregulated pathways. In this manner, we can detect pathways which play a prominent role in tumorigenesis and pinpoint specific interactions in the pathway that make a large contribution to its dysregulation and the disease phenotype.

We show that *dce* can recover significant DCEs and outperforms competitors in simulations. In a validation on real data we apply *dce* to a public CRISPR (Clustered Regularly Interspaced Short Palindromic Repeats) data set to recover differential effects in the network. In an exploratory study, we apply *dce* to breast cancer samples and compare the DCEs among different cancer stages. We identify dysregulated edges common across stages as well as stage-specific edges.

## 2 Results

In this section, we first show the performance of *dce* and its competitors on simulated data and a CRISPR data set. Then, we use *dce* for an exploratory analysis of breast cancer data from TCGA and show the progression of pathway dysregulation over different cancer stages.

### 2.1 Simulation study

Pathway databases contain networks of different sizes. Hence, we first investigate how much the size of the network influences performance. *dce* achieves the highest accuracy for all three network sizes considered (50, 100, 150 genes). Methods which do not account for confounding variables perform similar to random guessing for large networks (fig. 2a). However, *dce* also outperforms *pcor* with an AUC of 0.62 versus 0.56.

**Figure 2:**
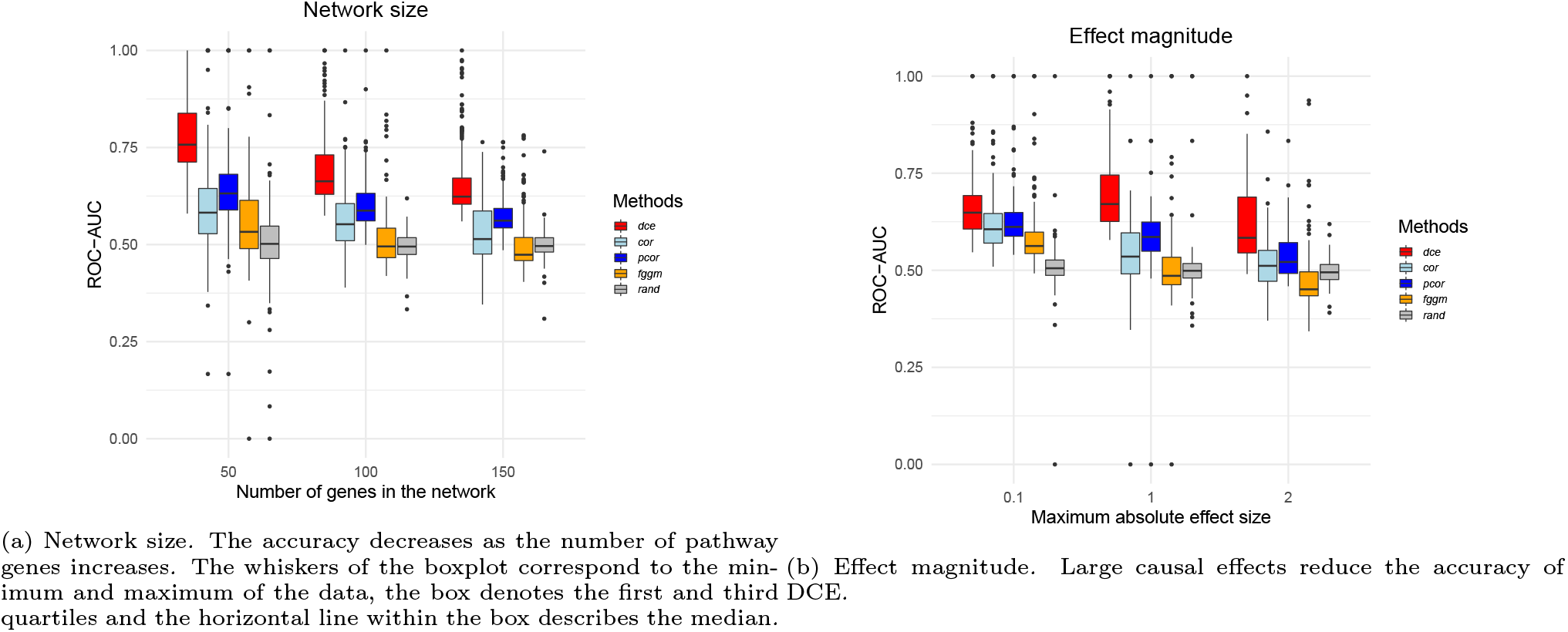
Performance benchmark. *dce* is compared to several competitors for varying network size (a) and effect magnitude (b) over 100 synthetic data sets each. *dce* achieves the highest accuracy, which decreases for large networks 𝒢 and very large or small differential effects. The whiskers of the boxplot correspond to the minimum and maximum of the data, the box denotes the first and third quartiles and the horizontal line within the box describes the median.

Second, we assess how the size of the effect magnitude affects the identification of significant differences. We varied the effect magnitude in the set {0.1, 1, 2}. For example, for an effect magnitude of 1 the edge weights between the network of the wild type samples and the disease samples differ by at most 1. *dce* has difficulty estimating large differences as well as very small differences. However, it still significantly outperforms all other methods, which again show similar performance to random guessing for large effects (fig. 2b).

In additional simulations, *dce* shows increasing accuracy for decreasing dispersion and increasing number of samples (figs. S1 and S2) as is expected due to decreasing noise. We found constant accuracy of *dce* over varying ranges of library size (fig. S3). Different prevalence of positive edges has little effect on the accuracy of *dce* (fig. S4). *dce* with latent variable adjustment performs similarly to *dce* without latent variable integration if we do not simulate any latent variables. But *dce* significantly outperforms *dce* without latent variable integration for five and ten latent variables influencing the data set (fig. S5). This is because without latent confounding adjustment one has a large number of false positives due to the confounding bias (fig S6).

*dce* relies heavily on the given network 𝒢. Hence, we investigate how well *dce* performs if 𝒢 contains false edges or is missing true edges. We find that *dce* is robust to additional false edges in the network, but starts breaking down if true edges are missing in larger fractions (fig. S7).

### 2.2 Validation experiments using CRISPR knockout data

To benchmark our method using real-life data generated by Perturb-seq [23], we ask whether we can recover the CRISPR knockout from single-cell RNA-seq data using the pathway “Protein processing in endoplasmic reticulum” (hsa04141) from KEGG. Hence we assume that hsa04141 captures the causal gene interactions governing the response of the cell to the experimental intervention. As seen in the synthetic benchmark, slight deviations of the observed network from the true underlying network have no major impact on the performance of our method (fig. S7). We then run *dce* on the single-cell gene expression data for each of the 14 conditions independently. By interpreting a CRISPR knockout as an intervention of the causal pathway, we define the positive class to consist of all edges adjacent to a knocked-out gene, and the negative class as all other genes. Consequently, a true positive occurs when an edge adjacent to a CRISPR knocked-out gene is (significantly) associated to a non-zero DCE.

Figure 3a shows an example of this procedure for one of the conditions described above. The CRISPR knockout gene is highlighted in red and a positive DCE of ∼ 1.3 can be observed on the edge connecting *ATF6* and *DDIT3*. This can be seen in more detail in fig. 3b. As this edge is adjacent to the knocked out gene *ATF6*, it is classified as a true positive for an effect size threshold of |0.5|. Following an analogous argument, the edge from *EIF2AK3* to *EIF2S1* is classified as a false positive.

**Figure 3:**
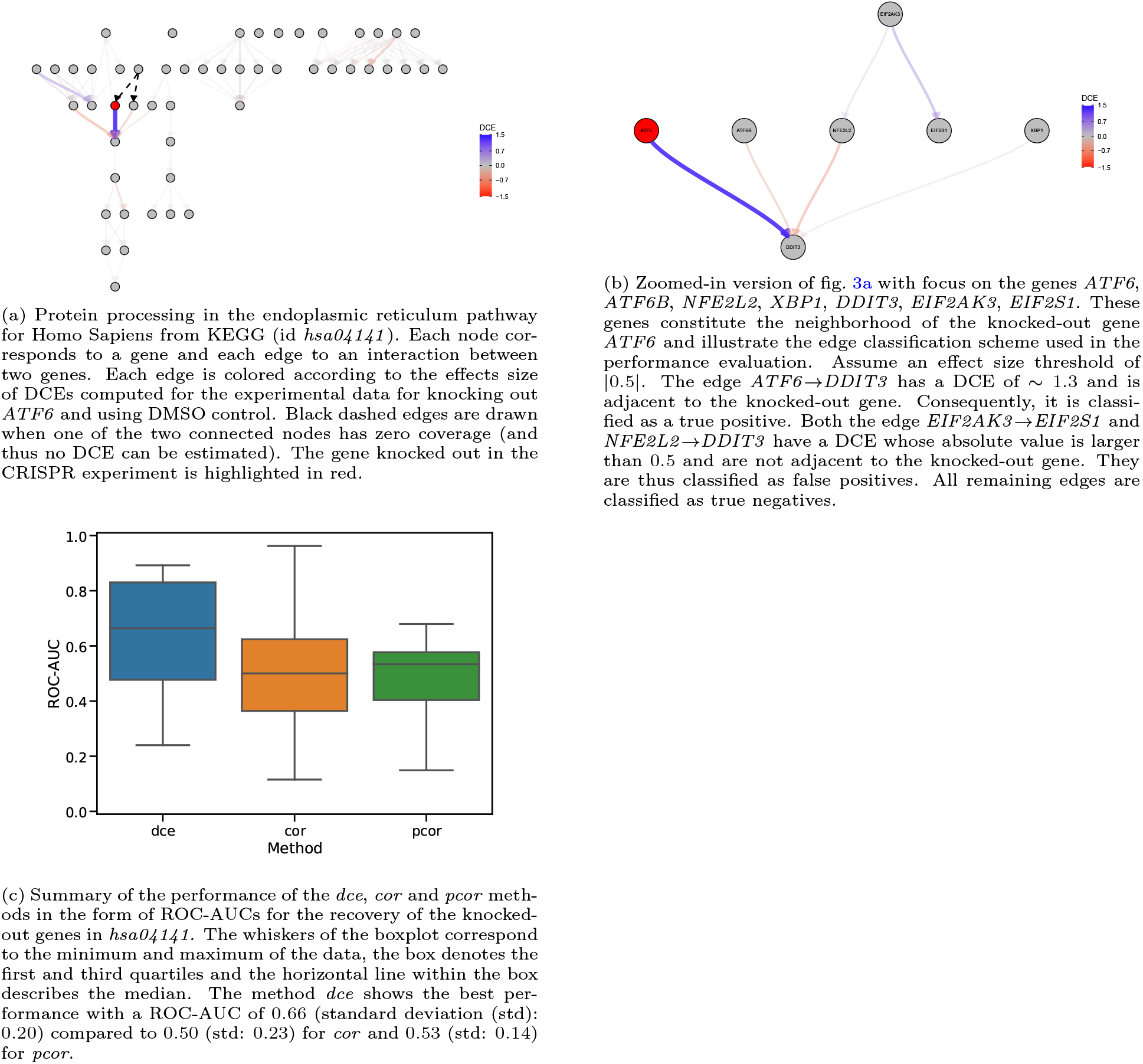
Overview of the CRISPR benchmark.

We find that *dce* better recovers the knock-out effect with a median ROC-AUC of 0.66 compared to 0.50 for *cor* and 0.53 for *pcor* (fig. 3c). This shows that *dce* is able to better recover the dysregulations of single as well as combinatorial knock-outs when compared to methods based on correlations.

### 2.3 Exploratory analysis of TCGA data

To demonstrate the ability of our method to recover known cancer-related pathway dysregulations as well as to discover new genes of potential biological and clinical relevance, we compute DCEs using breast cancer gene expression data from TCGA on the breast cancer pathway obtained from KEGG. The results for each stage are then visualized on the pathway structure (figs. 4a to 4c). The raw DCE values were transformed to a symmetric logarithm for greater visibility with the following formula

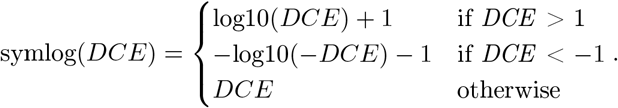

**Figure 4:**
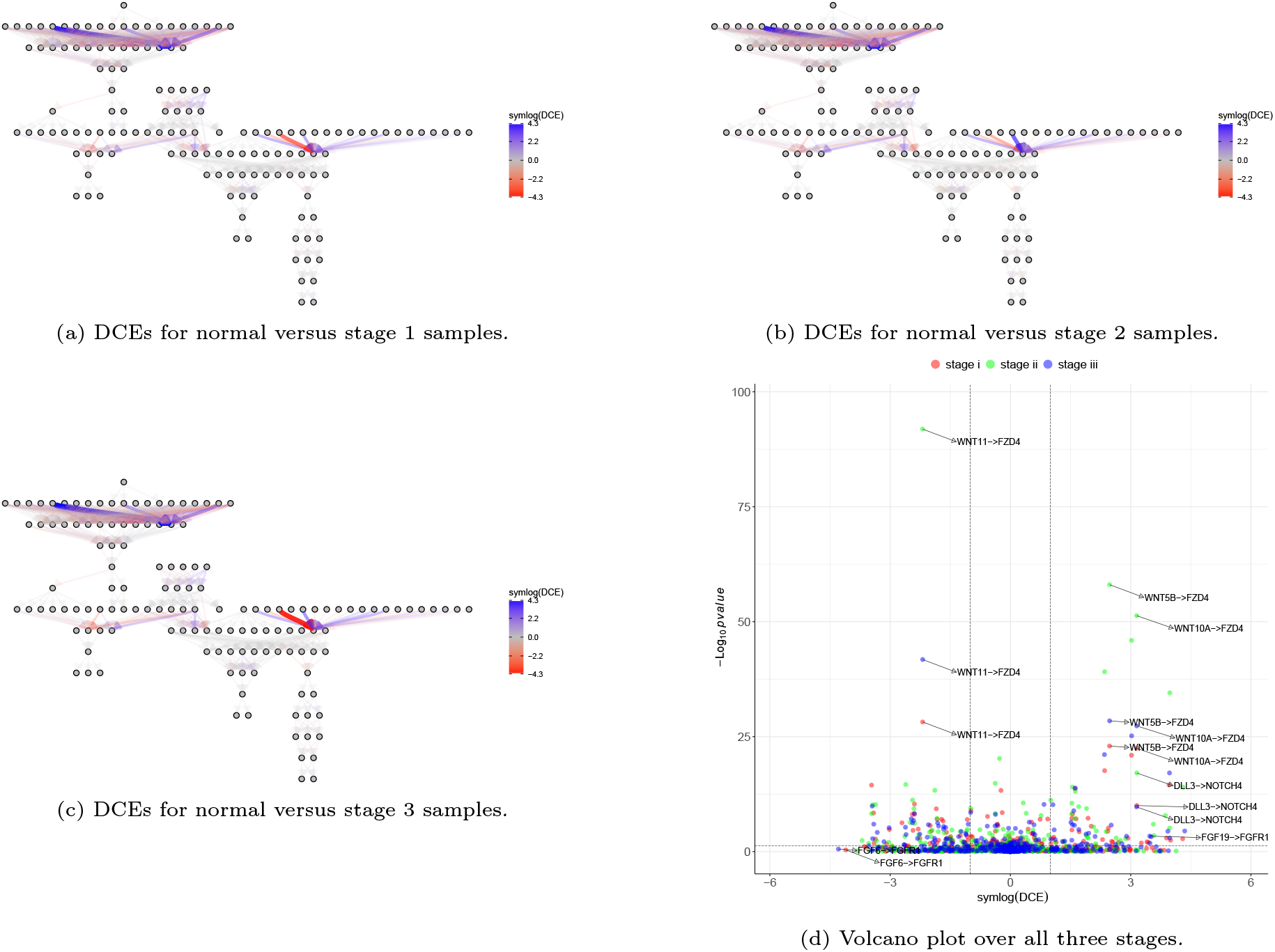
DCEs for *TCGA-BRCA normal* samples versus *stage I, stage II*, and *stage III* computed with the *hsa05224* pathway. In (a)-(c), edge thickness and opacity scale with absolute DCE size. More negative DCEs appear red, more positive DCEs appear blue. The color follows a symmetric logarithmic scale for values |*x*| *≥* 1 and is linear otherwise. (d) shows a volcano plot for the symmetric logarithm of DCE against its associated p-value. DCE thresholds of 1 and *−*1 as well as as p-value threshold of 0.05 are denoted with grey dashed lines.

Most edges do not show any difference in causal effect strength between normal and stage condition in all stages (fig. 4d). Throughout all stages, interactions between the WNT and FZD protein complexes exhibit significantly non-zero DCEs indicating a strong dysregulation of the breast cancer pathway. This pair has been implicated in disease formation in general [24–26] and in breast cancer in particular [27, 28]. We also observe a significant dysregulation of the DLL3 → NOTCH4 edge in all three stages. The Notch signaling pathway has been shown to play an important role in Pancreatic ductal adenocarcinoma tumor cells suggesting that stromal cells located in the breast may play an important role [29]. Though not significant (p-value *>* 0.05), interactions between the FGF and FGFR protein complexes show strong negative effect sizes in all three stages (DCE *< −* 100 for most members of these complexes). This pair has already been recognized as a promising therapeutic target for breast cancer treatment [30].

## 3 Discussion

We have presented a new method, *dce*, to compute differential causal effects between two conditions using a regression approach. *dce* enables the edge-specific identification of signaling pathway dysregulations. This piece of information can help to further our understanding of subtle differences on the molecular level in seemingly similar cancer types.

*dce* assumes a linear relationship among pathway genes. The linear model is solved using network information to account for additional genes confounding the linear relationship between gene pairs. The network information is included via prior knowledge from literature. *dce* also includes latent variables in the model accounting, e.g., for batch effects, which are unknown and not included in the gene network, as confounders.

Our simulation study emphasizes the need for sufficiently large datasets when dealing with large pathways. At the same time we have shown that *dce* is able to detect changes in causal effects even in the presence of noise and for certain ranges of effect sizes. For a wide array of parameter choices, *dce* outperforms methods using (partial) correlation and *fggm*. Especially in the case of latent confounders we showed that *dce* with the integration of latent variables outperforms *dce* without, except if no latent confounders were used to simulate the data. In this case both methods are equally accurate. Hence, we recommend the integration of latent variables in the model as the default configuration.

In addition to the synthetic benchmark, we have also validated our method on real data derived from Perturb-seq experiments. We have shown that *dce* is able to recover the experimental knock-outs with better performance than correlations and partial correlations.

For breast cancer, we have shown that not all parts of the signaling pathway are perturbed and characteristic hotspots exist. Some causal effects between two genes are invariant to stage information, while other causal effects can vary in either magnitude or even sign of their effect size. This indicates that certain areas of such pathways are more relevant than others. This phenomenon has also been observed in other studies ([31, 32]). Some parts of a pathway seem to be either more conserved or just not relevant for tumorigenesis. This provides exciting opportunities to identify drugs which target certain parts of a pathway and might explain their efficacy. However, the robustness of our method depends on the availability of enough samples. In many cases, few are available and make our approach infeasible. While *dce* performs still better than random for even 10 samples, it is significantly worse than for higher sample sizes.

In summary, we have proposed a novel application of the concept of differential causal effects which describe the difference in causal effects between two conditions and developed a regression approach to compute those differences. We demonstrate their robustness in a simulation study, and point out interesting results in application to real data, e.g., we show that some dysregulated edges are consistent among breast cancer tumor stages I-III, but that other dysregulations are unique to each stage.

Future research should focus on modifying the regression to make working with small data sets more robust, for example, enforcing sparsity by the introduction of *L*_1_- or *L*_2_-norms on the coefficients to avoid outliers produced by artifacts in the data.

We have shown the performance of *dce* on count data from simulations and (single cell) RNA-Seq. However, *dce* is also suited to analyze other types of data like Gaussian data from log normal microarray intensities.

## 4 Methods

In this section, we describe the Differential Causal Effects (*dce*) method. We briefly review the causality framework and then introduce the model and computation of DCEs, including under potential latent confounding. We provide implementation details for obtaining both the estimates and their significance levels. Finally, we describe the generating mechanism for synthetic data used throughout the paper.

### Causality of biological pathways

First, we give a quick review of causality in the context of biological pathways. A gene pathway can be represented as a structural equation model (SEM) consisting of a directed acyclic graph (DAG) 𝒢 with nodes 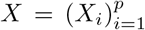 describing the expression of genes, a set of directed edges 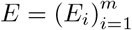 representing the causal structure and the structural equations 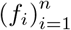 describing how each variable *X*_*i*_ is generated from its parents *X*_*pa*(*i*)_ in 𝒢, *X*_*i*_ ← *f*_*i*_(*X*_*pa*(*i*)_, *ϵ*_*i*_), where 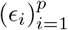 are jointly independent noise variables. The causal interpretation of an edge between any two nodes is as follows: changing the expression of a parent *X*_*i*_ affects the expression of the child node *X*_*j*_, which is propagated further to all descendants. The parental sets are given by the edge set *E*. Of particular interest are the interventional distributions for the SEM, in particular their expectations 𝔼 [*X*_*i*_ | *do*(*X*_*j*_ = *x*)], which describe how the expected value of the variable *X*_*i*_ changes when we intervene and set the variable *X*_*j*_ to some fixed value *x*. We define the causal effect (CE) of a variable *X*_*j*_ on its descendant *X*_*i*_ as

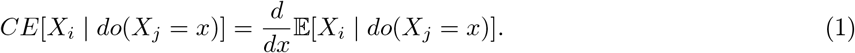

This derivative equals *β*_*x*_ if, by changing the value of *X*_*j*_ from *x* to *x* + Δ*x*, for some small value Δ*x*, the value of *X*_*i*_ changes on average by *β*_*x*_ Δ*x*. In the literature, the CE is often also referred to as the total causal effect, because it quantifies the overall effect of an intervention at variable *X*_*j*_ on all of its descendants. We are interested in differential causal effects (DCE) defined as the differences between the causal effects of two conditions of interest, such as, e.g., two different cancer stages or healthy and cancerous samples.

### Linearity of the conditional mean

We model the relationship between the mean of any gene expression *X*_*i*_ and its parents *X*_*pa*(*i*)_ by a linear function:

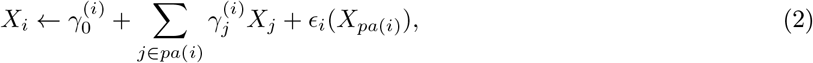

where, conditionally on *X*_*pa*(*i*)_, the error term ϵ _*i*_(*X*_*pa*(*i*)_) has mean zero, variance depending on *X*_*pa*(*i*)_ and 𝔼 [ϵ _*i*_(*X*_*pa*(*i*)_)|*X*_*pa*(*i*)_] ≡ 0. A prime example is any generalized linear model (GLM) with identity link function.

The coefficients 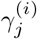 correspond to the direct causal effects, whereas the total causal effects (1) measure the aggregate effect over all directed paths from a certain variable *X*_*j*_ to *X*_*i*_ in 𝒢.

Let us consider two arbitrary genes *X*_*i*_ and *X*_*j*_ in the pathway. Under the linearity assumption, we can compute the causal effect *CE*[*X*_*i*_ | *do*(*X*_*j*_ = *x*)], which does not depend on *x*, as the coefficient *β* of the linear regression of *X*_*i*_ on *X*_*j*_ and an adjustment set 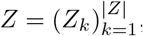,

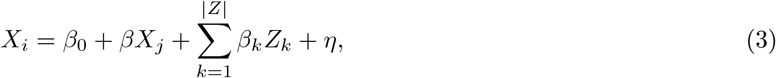

where *β*_0_ denotes the intercept and *η* is random noise with mean zero [33, 34]. The adjustment set *Z* is a set of nodes in the pathway 𝒢 which fulfills the Back-door criterion [7]. Hence, it holds that no element of *Z* is a descendant of *X*_*j*_, and *Z* blocks every path between *X*_*i*_ and *X*_*j*_ that contains an edge with *X*_*j*_ as the child. For example, the parent set *X*_*pa*(*j*)_ always fulfills the Back-door criterion and we always use it as adjustment set.

If the causal effects of the gene expression *X*_*j*_ on the gene expression *X*_*i*_ are denoted as *β*^*A*^ and *β*^*B*^ under different conditions *A* and *B*, respectively, then the differential causal effect (DCE) *δ* is obtained as the difference

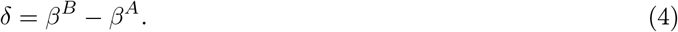

Given a graph 𝒢 describing a biological pathway and observations of the variables, we can compute all differential causal effects and identify interactions between any such two variables *X*_*j*_ and *X*_*i*_ that are different between the two conditions (fig. 1).

### Coupling the two conditions

We can compute the DCE *δ* on the edge *X*_*j*_ → *X*_*i*_ by fitting a joint model for both conditions. Let *I* be an indicator random variable, which is equal to 1, if the observation comes from condition A, and 0, if it comes from condition *B*. The DCE *δ* can be computed from all samples jointly by fitting the following linear model

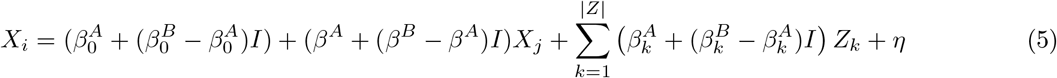

with interaction terms *IX*_*j*_ and *IZ*_*i*_. The differential causal effect *δ* = *β*^*B*^ *− β*^*A*^ can be estimated by using the coefficient estimate corresponding to the interaction term *IX*_*j*_ in (5).

### Testing for significance

Another advantage of fitting the joint model (5) over both conditions is that testing the significance of the estimated DCEs corresponds to the well-known task of testing the significance of coefficient estimates in a linear model. However, some care is needed if the variances of the error terms ϵ_*i*_(*X*_*pa*(*i*)_) in our structural equations (2) indeed depend on the values of the predictors *X*_*pa*(*i*)_, i.e., if there is a certain mean-variance relationship for the gene expression levels, as has been described for RNA-seq data [35]. In this case, the linear model (5) is heteroscedastic and the usual formulas for standard errors of the coefficient estimates, that result in t-tests for the significance, do not apply. We therefore use heteroscedasticity-consistent standard errors that yield asymptotically valid confidence intervals and p-values regardless of the dependence of the noise level on predictor values [36–38].

Besides assessing significance of DCE for single edges, we can also calculate a global p-value measuring the overall dysregulation of a given pathway 𝒢: we combine the p-values corresponding to different differential causal effects 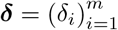 by taking their harmonic mean [39].

### Adjusting for latent confounding

A fundamental assumption in most of the causal inference methods is that there is no unobserved confounding, i.e., that there are no factors affecting both the cause and the effect [40, 41]. For example, batch effects due to varying laboratory conditions could act as such unobserved confounders. Presence of latent confounding can result in spurious correlations and false causal conclusions. Therefore, adjusting for potential latent confounding is crucial for making the method robust in applications to biological data [42].

Some information about latent factors can often be obtained from the principal components of the data [43]. This can be made rigorous under the linearity assumption (2), for our structural equation model 𝒢, as follows. We assume that there are *q* latent variables *H*_1_, …, *H*_*q*_ affecting our data. We extend the model (2) to include the latent confounding as follows:

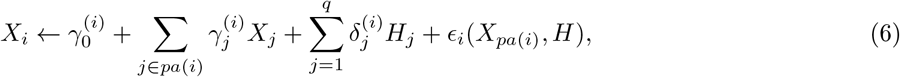

i.e., the latent confounders *H*_1_, …, *H*_*q*_ are additional source nodes in the DAG 𝒢 and affect every gene linearly, analogously to (2). By writing the structural equations (6) in matrix form with 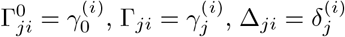 and *E*(*X, H*)_*ji*_ = ϵ _*i*_(*X*_*pa*(*i*)_, *H*)_*j*_, we obtain

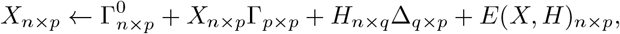

which gives

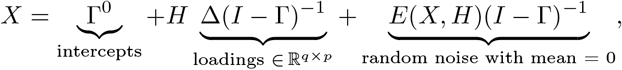

which is the standard linear factor model with heteroscedastic errors. From this representation, one can see that *H* can be determined from the principal components of *X* (fig. 5). The screeplot for a toy example visualizes the effect of latent variables having a global effect on the data. The first principle components are clearly separated from the rest, if latent factors are present (fig. 5, left). Therefore we obtain the confounding proxies *Ĥ* as the scores of the first 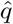 principal components of the design matrix combining the data from both conditions.

**Figure 5:**
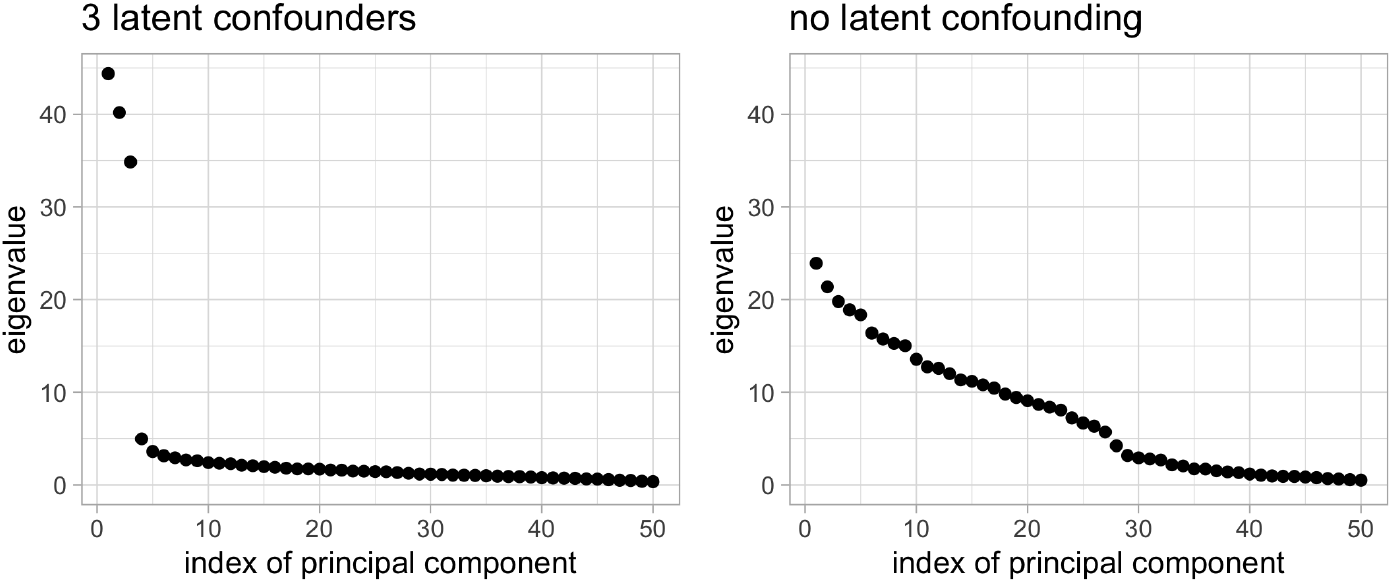
The screeplot (of synthetic data generated as described in the Methods section) shows that in presence of latent confounding as in (6), the first *q* principal components explain much more variability of the data, which we exploit for confounding adjustment.

The confounding proxies *Ĥ* are then simply added to the adjustment set *Z*, see equations (3) and (5). In this way, the Back-door adjustment not only adjusts for the confounding variables observed in the DAG 𝒢 as before, but also helps reducing the bias induced by latent confounding.

The deconfounding methodology relies on the assumption that every confounding variable affects many variables in the dataset, i.e., the confounding is dense [44]. In this case, we have a lot of information about the latent factors in the data and the confounding proxies *Ĥ* capture the effect of the confounders *H* well. Furthermore, the dense confounding assumption ensures that the scree plot, showing the singular values of the design matrix, has a spiked structure, as several latent factors can explain a relatively large proportion of the variance (fig. 5). This helps estimating the number 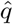 of the confounding proxies used. As a default choice, we use a permutation method that can be shown to work well under certain assumptions [45] and which compares the observed value of the variance explained by the principal components with its expected value over many random permutations of the values in each column of gene expression matrix *X*.

### Algorithm and implementation in R

The presented methods are implemented in the R package *dce* which is freely available on Bioconductor. The function *dce::dce* takes as input the structure of a biological pathway, i.e., the adjacency matrix of a DAG, and two *n* × *p* matrices, with *n* samples and *p* genes, storing gene expression data for each of the two conditions respectively. As output, the function returns the estimated DCEs, as well as standard errors and two-sided p-values for the DCE at each edge in the pathway. The results can be easily transformed into a dataframe and plotted for further downstream analyses, together with the p-value measuring the overall pathway enrichment.

### Generating synthetic data and benchmarking methods

We assess the behavior of *dce* and its competitors in a controlled setting by generating synthetic data with known DCEs (ground truth). We start by generating a random DAG 𝒢. Without loss of generality, we assume the nodes of the DAG to be topologically ordered, i.e., node *X*_*i*_ can only be parent of node *X*_*j*_, if *i < j*. This ensures that the network 𝒢 is a DAG. In practice, we sample edges from a binomial distribution with probability 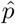 for the upper triangle of 𝒢. We further sample the coefficients 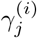 for every edge as in (2) from a uniform distribution 𝒰 (−*γ*_max_, *γ*_max_). We generate the data for network 𝒢 in the following way. For a node *X*_*i*_, we set the mean expression count

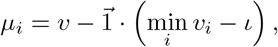

where each *X*_*j*_ is a vector of counts, corresponding to gene expression values from experiments like RNA-Seq. 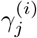 represents the direct effect of *X*_*j*_ on *X*_*i*_, 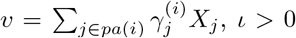 is a small shift, and 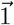 is a vector of ones. Subtracting the minimum ensures positive values of the mean for each data point. Then, a realization of *X*_*i*_ is drawn from the Poisson distribution Pois 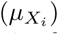. We introduce negative binomial noise by drawing a realization of each source node in 𝒢 from the negative binomial distribution NB (*µ, θ*) with a general mean *µ* and dispersion *θ*. We use this setup to control the variance across all nodes, which can blow up for descendants with larger means.

After sampling the data *D*_*A*_ for the nodes of network 𝒢 under condition *A*, we resample a certain fraction of edge weights in order to generate new data *D*_*B*_ under condition *B*. For an edge weight *β*^*A*^ we sample the new edge weight from a uniform distribution

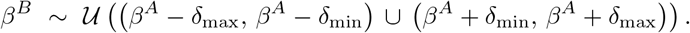

This ensures that the absolute difference between the two edge weights lies in [*δ*_min_, *δ*_max_].

We also simulate latent variables. They are neither included in the data nor the network 𝒢, but have (unknown) outgoing edges to all genes in the data set with non-zero effects. Hence, these latent variables have global effects on the data, e.g., emulating batch effects.

We compare *dce* it to correlation (*cor*), partial correlation (*pcor*), the method Fast Gaussian Graphical Models (*fggm*) tailored to DCEs [17, 46], and random guessing. *fggm* is based on partial correlation, but additionally tries to learn the network structure to adjust for confounding effects. We provide *pcor* with the same adjustment set of confounding variables as *dce*. We run all methods on simulated data for various modeling parameters. The default parameters are a network 𝒢 of 100 genes, 200 samples for both sample conditions, an absolute magnitude in effect differences between the two conditions of 1, mean of 100 negative binomial distributed counts with a dispersion of 1 for the source genes in the network 𝒢 (no parents), a true positive rate of 50% (edges which have different effects between the two conditions), and library size factors for each sample in the interval [1, 10]. The library size factor accounts for different sequencing depth among the samples, i.e., for one sample including more reads because more RNA was available even though the gene expression was the same as in samples with less RNA. We account for different library sizes over all samples by computing transcripts per million (tpm).

Overall we simulate a full data set of 10, 000 genes including the genes in the network 𝒢 to allow for the realistic estimation of the library size. As a performance measure we use the area under the receiver operating characteristic (ROC-AUC). We count the number of true/false positive and false negative DCEs based on the edges in the ground truth network and the significant p-values for different significance levels. Based on these true/false positives we can compute the ROC curve and its AUC. For both correlation methods we use a permutation test to compute empirical p-values.

### Validation using Perturb-seq

Perturb-seq, a CRISPR-Cas9-based gene knockout method, can be used to inhibit the expression of multiple target genes on a single-cell level [23, 47]. In [23], this approach was used to systematically analyze the response of an integrated endoplasmic reticulum (ER) stress response pathway to the combinatorial knockout of the three transmembrane sensor proteins IRE1*α*, ATF6, and PERK. Each combinatorial knockout was treated either with a DMSO control, tunicamycin, or thapsigargin. In total, this yielded 21 conditions. In the following analysis, we used the data sets for transduced cells and excluded the thapsigargin treatment due to very low coverage (median coverage of knocked-out gene and its neighborhood for the three treatments: DMSO control: 1.9, tunicamycin: 2.3, thapsigargin: 0.9). We downloaded the raw data from NCBI GEO (accession: GSE90546) and mapped guide RNA barcodes to cell barcodes to create a gene expression count matrix of the individual cells labeled by their corresponding knock-outs. We compare the performance of *dce* to *cor* and *pcor*. We estimate the significance of whether a difference in correlation is different from zero using a permutation test. The performance of each method is evaluated using the area-under-curve (AUC) metric for the receiver-operating-characteristic (ROC) curve. The false and true positive rates for the ROC curve are computed from the p-value per edge as in the synthetic benchmark.

### Exploratory analysis with TCGA data

We retrieve gene expression data from The Cancer Genome Atlas (TCGA) [48] and pathway structures from the Kyoto Encyclopedia of Genes and Genomes (KEGG) [11]. Unlike the Perturb-seq dataset, data obtained from TCGA is observational instead of interventional. We do thus not have any ground truth information and perform an exploratory analysis. For a given cancer type, the associated samples are first grouped into normal and tumor samples. The tumor samples are subsequently stratified according to their stage. In particular, we downloaded all normal and tumor gene expression samples from TCGA for breast cancer (*TCGA-BRCA*) and selected all stages with a sufficient number of samples (*stage I*: 202 samples, *stage II*: 697 samples, *stage III*: 276 samples; *normal*: 113 samples). We use the breast cancer pathway (*hsa05224*) from KEGG which contains 147 nodes and 509 edges. We then compute DCEs between the normal condition and each of the three stages of the tumor condition, respectively.

## 5 Data availability

The code used to construct the synthetic data sets is available as part of the R software package *dce*. The experimental data used in the Perturb-seq validation is available under the accession GSE90546 from NCBI GEO. The experimental data used in the exploratory breast cancer analysis is available under the accession TCGA-BRCA from The Cancer Genome Atlas. The pathway structures have been obtained from the Kyoto Encyclopedia of Genes and Genome.

## 6 Code availability

The method *dce* is freely available as an R package on Bioconductor as well as on https://github.com/cbg-ethz/dce. The GitHub repository also contains the Snakemake [49] workflows needed to reproduce all results presented here.

## 7 Acknowledgements

Part of this work has been funded by SystemsX.ch, the Swiss Initiative in Systems Biology, under Grant No. RTD 2013/152 (TargetInfectX - Multi-Pronged Perturbation of Pathogen Infection in Human Cells), evaluated by the Swiss National Science Foundation, and by ERC Synergy Grant 609883 (to NB). The research of DC and PB has been supported by the European Research Council under the Grant Agreement No. 786461 (CausalStats - ERC-2017-ADG).

## 8 Author information

### Contributions

KPJ and MP conceived the project. KPJ and MP developed the statistical model of *dce* and implemented the software package. DC contributed to the statistical methodology as well as software implementation. NB and PB supervised the study. KPJ and MP wrote the initial manuscript draft. All authors edited the manuscript.

## References

[1] “The global challenge of cancer”, Nature Cancer, vol. 1, no. 1, pp. 1–2, 2020. doi: 10.1038/s43018-019-0023-9. [Online]. Available: https://doi.org/10.1038/s43018-019-0023-9.

[2] N. Hawkes, “Cancer survival data emphasise importance of early diagnosis”, BMJ, vol. 364, 2019, issn: 0959-8138. doi: 10.1136/bmj.l408. eprint: https://www.bmj.com/content/364/bmj.l408.full.pdf. [Online]. Available: https://www.bmj.com/content/364/bmj.l408.

[3] K. D. Miller, L. Nogueira, A. B. Mariotto, J. H. Rowland, K. R. Yabroff, C. M. Alfano, A. Jemal, J. L. Kramer, and R. L. Siegel, “Cancer treatment and survivorship statistics, 2019”, CA: a cancer journal for clinicians, vol. 69, no. 5, pp. 363–385, 2019.

[4] M. A. Troester and T. Swift-Scanlan, “Challenges in studying the etiology of breast cancer subtypes”, Breast Cancer Research, vol. 11, no. 3, p. 104, 2009. doi: 10.1186/bcr2323. [Online]. Available: https://doi.org/10.1186/bcr2323.

[5] S. Khakabimamaghani, D. Ding, O. Snow, and M. Ester, “Uncovering the subtype-specific temporal order of cancer pathway dysregulation”, PLoS computational biology, vol. 15, no. 11, 2019.

[6] D. Hanahan and R. A. Weinberg, “Hallmarks of cancer: The next generation”, cell, vol. 144, no. 5, pp. 646– 674, 2011.

[7] J. Pearl, Causality: Models, Reasoning and Inference. Cambridge University Press, Cambridge, UK., 2000.

[8] B. Vogelstein, N. Papadopoulos, V. E. Velculescu, S. Zhou, L. A. Diaz, and K. W. Kinzler, “Cancer genome landscapes”, science, vol. 339, no. 6127, pp. 1546–1558, 2013.

[9] A. Subramanian, P. Tamayo, V. K. Mootha, S. Mukherjee, B. L. Ebert, M. A. Gillette, A. Paulovich, S. L. Pomeroy, T. R. Golub, E. S. Lander, et al., “Gene set enrichment analysis: A knowledge-based approach for interpreting genome-wide expression profiles”, Proceedings of the National Academy of Sciences, vol. 102, no. 43, pp. 15 545–15 550, 2005.

[10] D. Wu and G. K. Smyth, “Camera: A competitive gene set test accounting for inter-gene correlation”, Nucleic acids research, vol. 40, no. 17, e133–e133, 2012.

[11] H. Ogata, S. Goto, K. Sato, W. Fujibuchi, H. Bono, and M. Kanehisa, “Kegg: Kyoto encyclopedia of genes and genomes”, Nucleic acids research, vol. 27, no. 1, pp. 29–34, 1999.

[12] A. Liu, P. Trairatphisan, E. Gjerga, A. Didangelos, J. Barratt, and J. Saez-Rodriguez, “From expression footprints to causal pathways: Contextualizing large signaling networks with carnival”, NPJ systems biology and applications, vol. 5, no. 1, pp. 1–10, 2019.

[13] B. Dutta, A. Wallqvist, and J. Reifman, “Pathnet: A tool for pathway analysis using topological information”, Source code for biology and medicine, vol. 7, no. 1, p. 10, 2012.

[14] A. L. Tarca, S. Draghici, P. Khatri, S. S. Hassan, P. Mittal, J.-s. Kim, C. J. Kim, J. P. Kusanovic, and R. Romero, “A novel signaling pathway impact analysis”, Bioinformatics, vol. 25, no. 1, pp. 75–82, 2009.

[15] C. J. Vaske, S. C. Benz, J. Z. Sanborn, D. Earl, C. Szeto, J. Zhu, D. Haussler, and J. M. Stuart, “Inference of patient-specific pathway activities from multi-dimensional cancer genomics data using paradigm”, Bioinformatics, vol. 26, no. 12, pp. i237–i245, 2010.

[16] Y. Wang, C. Squires, A. Belyaeva, and C. Uhler, “Direct estimation of differences in causal graphs”, in Advances in Neural Information Processing Systems, 2018, pp. 3770–3781.

[17] H. He, S. Cao, J.-g. Zhang, H. Shen, Y.-P. Wang, and H.-w. Deng, “A statistical test for differential network analysis based on inference of gaussian graphical model”, Scientific Reports, vol. 9, no. 1, p. 10 863, 2019.

[18] D. Tian, Q. Gu, and J. Ma, “Identifying gene regulatory network rewiring using latent differential graphical models”, Nucleic acids research, vol. 44, no. 17, e140–e140, 2016.

[19] D. Nishimura, “Biocarta”, Biotech Software & Internet Report: The Computer Software Journal for Scient, vol. 2, no. 3, pp. 117–120, 2001.

[20] M. Whirl-Carrillo, E. McDonogh, J. Herbet, L. Gong, K. Sangkuhl, C. Thotn, R. Altman, and E. Klein, Pharmacogenomics knowledge for personlized medicine. clinical pharmacology and therpeutics 92, 4 (2012), 414–417, 2012.

[21] H. Mi, D. Ebert, A. Muruganujan, C. Mills, L.-P. Albou, T. Mushayamaha, and P. D. Thomas, “Panther version 16: A revised family classification, tree-based classification tool, enhancer regions and extensive api”, Nucleic Acids Research, vol. 49, no. D1, pp. D394–D403, 2021.

[22] C. F. Schaefer, K. Anthony, S. Krupa, J. Buchoff, M. Day, T. Hannay, and K. H. Buetow, “Pid: The pathway interaction database”, Nucleic acids research, vol. 37, no. Suppl_1, pp. D674–D679, 2009.

[23] B. Adamson, T. M. Norman, M. Jost, M. Y. Cho, J. K. Nuñez, Y. Chen, J. E. Villalta, L. A. Gilbert, M. A. Horlbeck, M. Y. Hein, et al., “A multiplexed single-cell crispr screening platform enables systematic dissection of the unfolded protein response”, Cell, vol. 167, no. 7, pp. 1867–1882, 2016.

[24] J. P. Dijksterhuis, B. Baljinnyam, K. Stanger, H. O. Sercan, Y. Ji, O. Andres, J. S. Rubin, R. N. Hannoush, and G. Schulte, “Systematic mapping of wnt-fzd protein interactions reveals functional selectivity by distinct wnt-fzd pairs”, Journal of Biological Chemistry, vol. 290, no. 11, pp. 6789–6798, 2015.

[25] A. J. Chien, W. H. Conrad, and R. T. Moon, “A wnt survival guide: From flies to human disease”, Journal of Investigative Dermatology, vol. 129, no. 7, pp. 1614–1627, 2009.

[26] G. Schulte, “International union of basic and clinical pharmacology. lxxx. the class frizzled receptors”, Pharmacological reviews, vol. 62, no. 4, pp. 632–667, 2010.

[27] P. Yin, W. Wang, J. Gao, Y. Bai, Z. Wang, L. Na, Y. Sun, and C. Zhao, “Fzd2 contributes to breast cancer cell mesenchymal-like stemness and drug resistance”, Oncology Research Featuring Preclinical and Clinical Cancer Therapeutics, vol. 28, no. 3, pp. 273–284, 2020.

[28] A. Koval and V. L. Katanaev, “Dramatic dysbalancing of the wnt pathway in breast cancers”, Scientific reports, vol. 8, no. 1, pp. 1–10, 2018.

[29] H. Song and Y. Zhang, “Regulation of pancreatic stellate cell activation by notch3”, BMC cancer, vol. 18, no. 1, pp. 1–12, 2018.

[30] M. F. Santolla and M. Maggiolini, “The fgf/fgfr system in breast cancer: Oncogenic features and therapeutic perspectives”, Cancers, vol. 12, no. 10, p. 3029, 2020.

[31] D. Song, M. Cui, G. Zhao, Z. Fan, K. Nolan, Y. Yang, P. Lee, F. Ye, and D. Y. Zhang, “Pathway-based analysis of breast cancer.”, eng, Am J Transl Res, vol. 6, no. 3, pp. 302–311, 2014, issn: 1943-8141 (Print); 1943-8141 (Electronic); 1943-8141 (Linking).

[32] Y. Feng, M. Spezia, S. Huang, C. Yuan, Z. Zeng, L. Zhang, X. Ji, W. Liu, B. Huang, W. Luo, B. Liu, Y. Lei, S. Du, A. Vuppalapati, H. H. Luu, R. C. Haydon, T.-C. He, and G. Ren, “Breast cancer development and progression: Risk factors, cancer stem cells, signaling pathways, genomics, and molecular pathogenesis”, Genes & diseases, vol. 5, no. 2, pp. 77–106, May 2018. doi: 10.1016/j.gendis.2018.05.001. [Online]. Available: https://pubmed.ncbi.nlm.nih.gov/30258937.

[33] M. Goldszmidt and J. Pearl, “Rank-based systems: A simple approach to belief revision, belief update, and reasoning about evidence and actions.”, in Proceeding of the 3rd Conference on Knowledge Representation, Jan. 1992, pp. 661–672.

[34] J. Pearl, “Causal diagrams for empirical research”, Biometrika, vol. 82, no. 4, pp. 669–688, 1995, issn: 00063444. [Online]. Available: http://www.jstor.org/stable/2337329.

[35] M. D. Robinson and G. K. Smyth, “Moderated statistical tests for assessing differences in tag abundance”, Bioinformatics, vol. 23, no. 21, pp. 2881–2887, Sep. 2007, issn: 1367-4803. doi: 10.1093/bioinformatics/btm453. eprint: https://academic.oup.com/bioinformatics/article-pdf/23/21/2881/16860418/btm453.pdf. [Online]. Available: https://doi.org/10.1093/bioinformatics/btm453.

[36] F. Eicker, “Limit theorems for regressions with unequal and dependent errors”, in Proceedings of the fifth Berkeley symposium on mathematical statistics and probability, Berkeley, CA: University of California Press, vol. 1, 1967, pp. 59–82.

[37] P. J. Huber et al., “The behavior of maximum likelihood estimates under nonstandard conditions”, in Proceedings of the fifth Berkeley symposium on mathematical statistics and probability, University of California Press, vol. 1, 1967, pp. 221–233.

[38] H. White, “A heteroskedasticity-consistent covariance matrix estimator and a direct test for heteroskedasticity”, Econometrica: journal of the Econometric Society, pp. 817–838, 1980.

[39] I. J. Good, “Significance tests in parallel and in series”, Journal of the American Statistical Association, vol. 53, no. 284, pp. 799–813, 1958.

[40] J. T. Leek, W. E. Johnson, H. S. Parker, A. E. Jaffe, and J. D. Storey, “The sva package for removing batch effects and other unwanted variation in high-throughput experiments”, Bioinformatics, vol. 28, no. 6, pp. 882–883, 2012.

[41] J. A. Gagnon-Bartsch, L. Jacob, and T. P. Speed, “Removing unwanted variation from high dimensional data with negative controls”, Berkeley: Tech Reports from Dep Stat Univ California, pp. 1–112, 2013.

[42] D. Ćevid, P. Bühlmann, and N. Meinshausen, “Spectral deconfounding via perturbed sparse linear models”, Journal of Machine Learning Research, vol. 21, p. 232, 2020.

[43] J. Novembre and M. Stephens, “Interpreting principal component analyses of spatial population genetic variation”, Nature genetics, vol. 40, no. 5, pp. 646–649, 2008.

[44] Z. Guo, D. Ćevid, and P. Bühlmann, “Doubly debiased lasso: High-dimensional inference under hidden confounding and measurement errors”, arXiv preprint 2004.03758, 2020.

[45] E. Dobriban, “Permutation methods for factor analysis and pca”, arXiv preprint 1710.00479, 2017.

[46] T. Wang, Z. Ren, Y. Ding, Z. Fang, Z. Sun, M. L. MacDonald, R. A. Sweet, J. Wang, and W. Chen, “Fastggm: An efficient algorithm for the inference of gaussian graphical model in biological networks”, PLOS Computational Biology, vol. 12, no. 2, pp. 1–16, Feb. 2016. doi: 10.1371/journal.pcbi.1004755. [Online]. Available: https://doi.org/10.1371/journal.pcbi.1004755.

[47] L. S. Qi, M. H. Larson, L. A. Gilbert, J. A. Doudna, J. S. Weissman, A. P. Arkin, and W. A. Lim, “Repurposing crispr as an rna-guided platform for sequence-specific control of gene expression”, Cell, vol. 152, no. 5, pp. 1173–1183, 2013.

[48] J. N. Weinstein, E. A. Collisson, G. B. Mills, K. R. M. Shaw, B. A. Ozenberger, K. Ellrott, I. Shmulevich, C. Sander, J. M. Stuart, C. G. A. R. Network, et al., “The cancer genome atlas pan-cancer analysis project”, Nature genetics, vol. 45, no. 10, p. 1113, 2013.

[49] F. Mölder, K. P. Jablonski, B. Letcher, M. B. Hall, C. H. Tomkins-Tinch, V. Sochat, J. Forster, S. Lee, S. O. Twardziok, A. Kanitz, et al., “Sustainable data analysis with snakemake”, F1000Research, vol. 10, no. 33, p. 33, 2021.

