## Supplementary figures for "Identifying cancer pathway dysregulations using differential causal effects"

1 Department of Biosystems Science and Engineering, ETH Zurich, Basel, 4058, Switzerland
2 SIB Swiss Institute of Bioinformatics, Basel, 4058, Switzerland
3 Seminar for Statistics, ETH Zürich, Zürich, 8092, Switzerland
* To whom correspondence should be addressed
$ Equal contribution

May 19, 2021

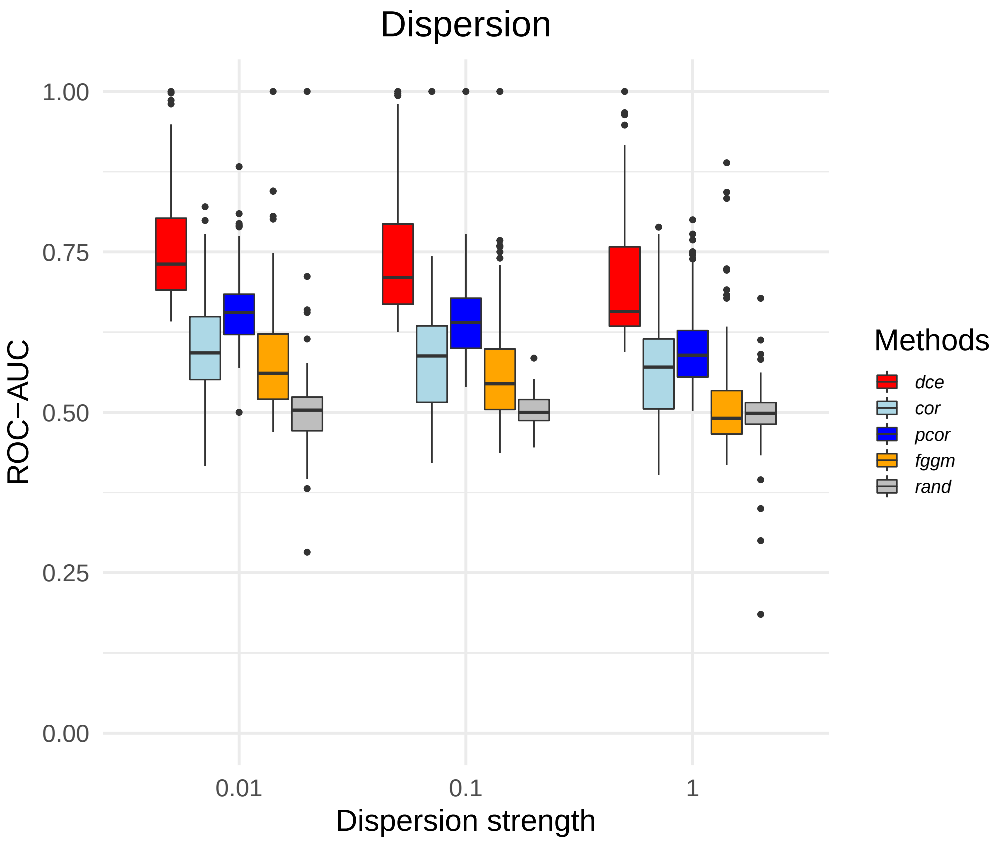


Figure S1: Dispersion. *dce* is compared to its competitors over 100 synthetic data sets with varying dispersion values. Performance decreases for higher dispersion values. The whiskers of the boxplot correspond to the minimum and maximum of the data, the box denotes the first and third quartiles and the horizontal line within the box describes the median.

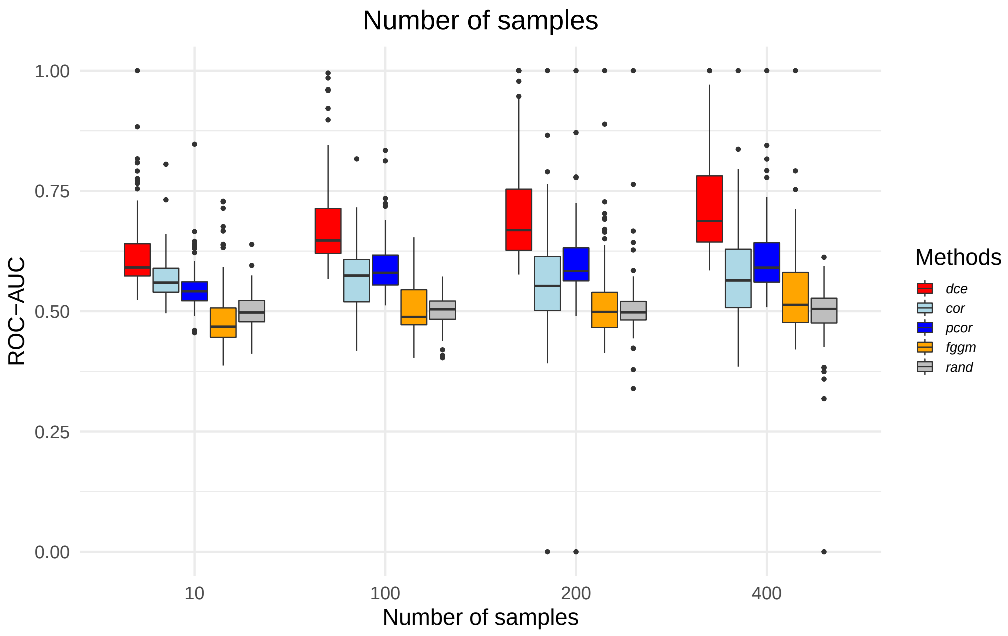


Figure S2: Sample size. *dce* is compared to its competitors over 100 synthetic data sets with varying sample sizes for one condition. The other conditions has a fixed sample size of 200. Performance decreases for lower sample sizes. The whiskers of the boxplot correspond to the minimum and maximum of the data, the box denotes the first and third quartiles and the horizontal line within the box describes the median.

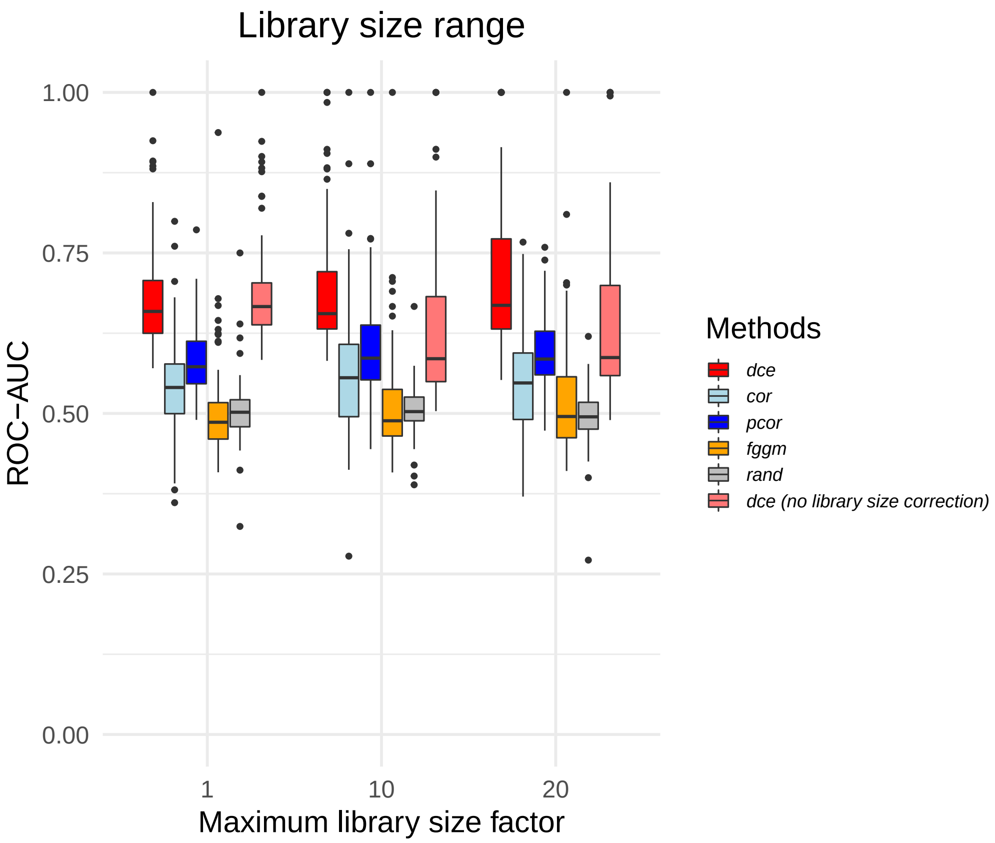


Figure S3: Library size. *dce* is compared to its competitors over 100 synthetic data sets with varying library size factors. Library size has little effect on the accuracy of all methods. The whiskers of the boxplot correspond to the minimum and maximum of the data, the box denotes the first and third quartiles and the horizontal line within the box describes the median.

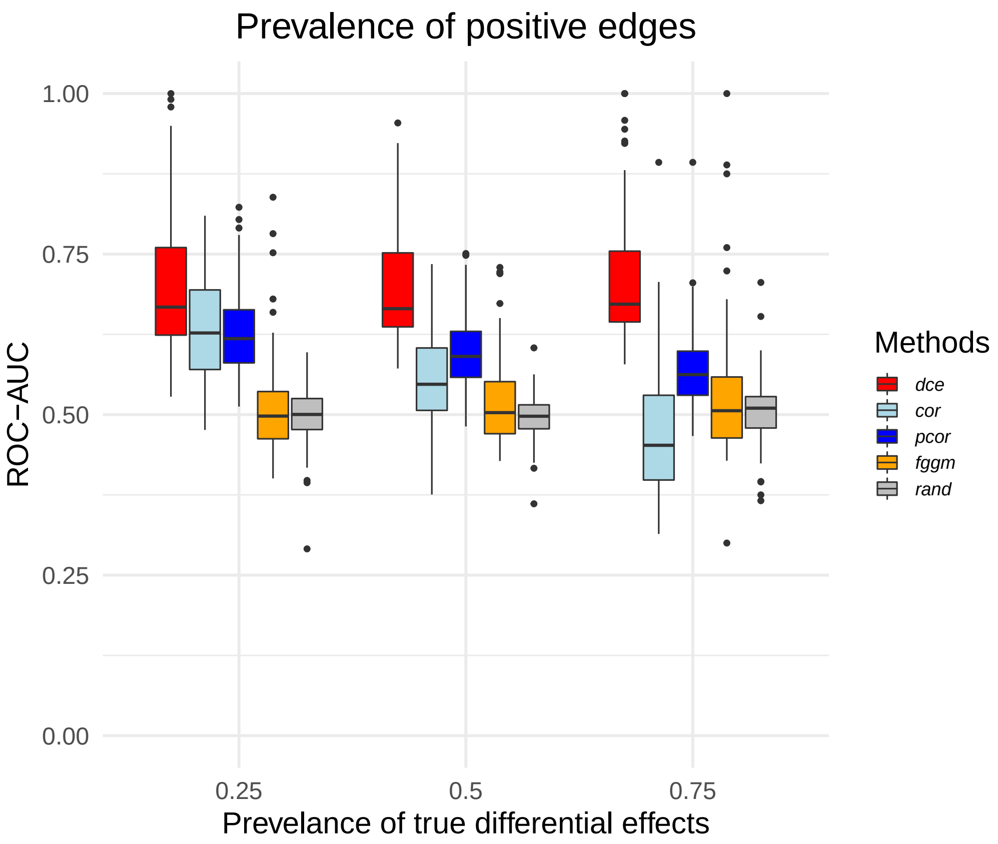


Figure S4: Prevalence. *dce* is compared to its competitors over 100 synthetic data sets with varying prevalence for DCE $\neq 0$. Accuracy decreases for all methods and higher prevalence except for *dce*. The whiskers of the boxplot correspond to the minimum and maximum of the data, the box denotes the first and third quartiles and the horizontal line within the box describes the median.

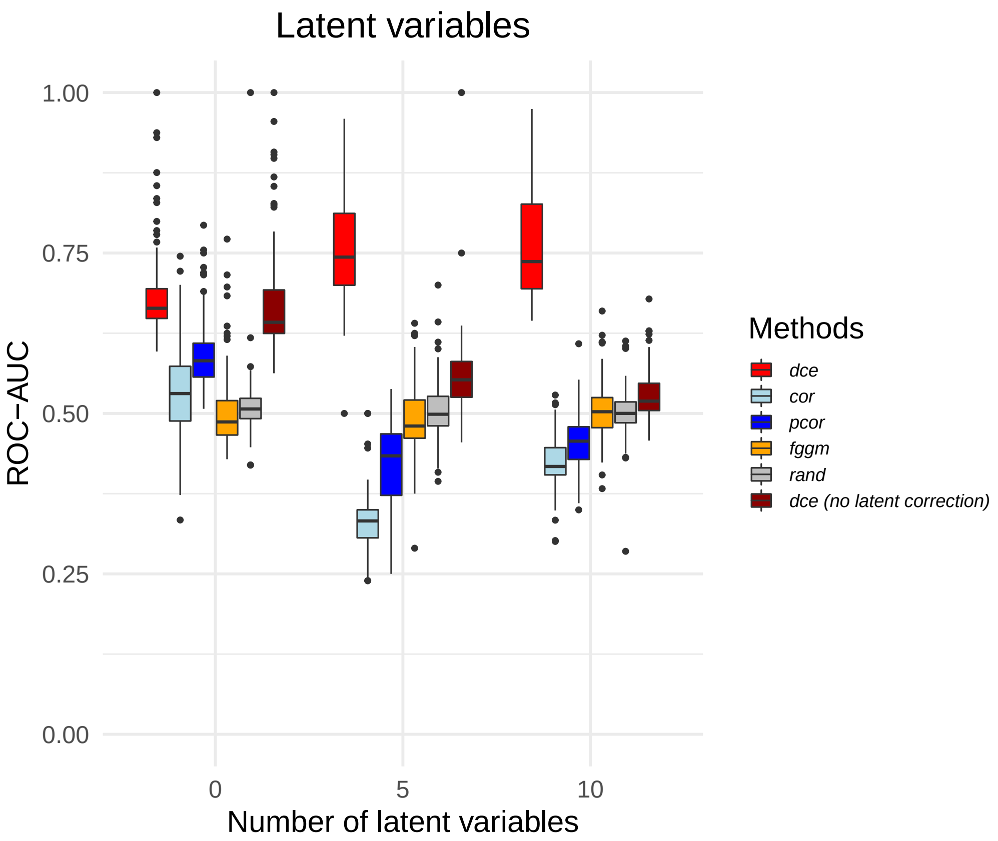


Figure S5: Latent variables. *dce* is compared to its competitors over 100 synthetic data sets with varying numbers of latent variables. *dce*'s accuracy stays robust, if we account for latent variables, but drastically decreases, if we do not. The whiskers of the boxplot correspond to the minimum and maximum of the data, the box denotes the first and third quartiles and the horizontal line within the box describes the median.

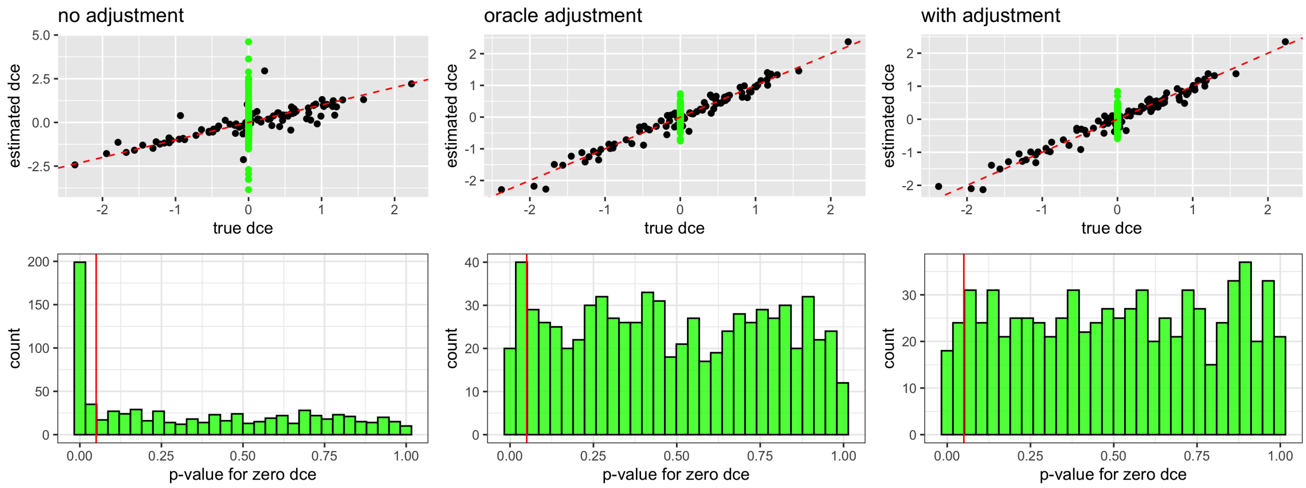


Figure S6: The performance of the *dce* without latent confounding adjustment (left), *dce* using true values of condounders (not known in practice) and *dce* with the latent confounding adjustment. Null DCEs are denoted in green, whereas the non-zero DCEs are denoted in black. This figure uses synthetic data with 300 genes, 300 observations and 3 latent confounders. Red line in the bottom row indicates the 0.05 threshold. The performance with the deconfounding step is close to the performance if we actually observed the latent confounders. Furthermore, it avoids increased number of falsely significant findings due to confounding bias (bottom row).

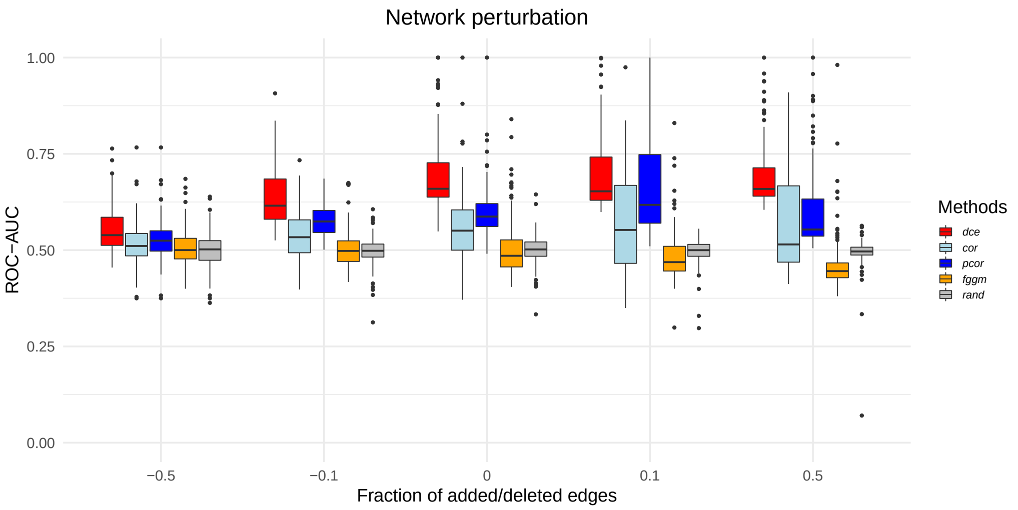


Figure S7: *dce* is compared to its competitors over 100 synthetic data sets with incorrect network information. Performance decreases for networks with missing edges, but stays robust, if additional edges are included. The whiskers of the boxplot correspond to the minimum and maximum of the data, the box denotes the first and third quartiles and the horizontal line within the box describes the median.
